# A Machine Learning Approach Elucidates Spatial Patterns of Environmental Properties Driving Microbial Composition Over Santos Basin, South Atlantic

**DOI:** 10.1101/2025.02.26.640418

**Authors:** Julio Cezar Fornazier Moreira, Flúvio Modolon, Natascha Menezes Bergo, Danilo Candido Vieira, Gustavo Fonseca, Francielli Vilela Peres, Rebeca Graciela Matheus Lizárraga, Diana Carolina Duque-Castaño, Alice de Moura Emilio, Augusto Miliorini Amendola, Renato Gamba Romano, Mateus Gustavo Chuqui, Fabiana da Silva Paula, Daniel Leite Moreira, Célio Roberto Jonck, Amanda Bendia, Frederico Pereira Brandini, Vivian Helena Pellizari

## Abstract

Microbial communities in marine ecosystems play a pivotal role in global biogeochemical cycles, with particular relevance in ecologically and industrially significant regions such as the Santos Basin (SB), Brazil’s largest marine sedimentary basin and a hub for oil and gas exploration. Yet, our capacity of predicting the microbial community structure and function remains limited for marine ecosystems. This study investigated the structure of microbial communities across different depths in the SB, using amplicon sequencing data and quantitative cell counts obtained via flow cytometry. Using a hybrid machine learning framework combining Self-Organizing Maps and Random Forest, we identified five distinct microbial assemblages (named microbial associations) in the SB predicted with 86% accuracy. These associations were primarily driven by temperature, water density, salinity, and nutrients such as phosphate and nitrate. Our findings showed a clear stratification of microbial communities across pelagic zones, with temperature as the main factor driving the structure in epipelagic and mesopelagic layers, while salinity and density exerted greater influence in the deeper bathypelagic communities. Temporal and spatial variations, particularly between 2019 and 2021, in regions influenced by the Cabo Frio upwelling and Rio de la Plata plume, further highlighted the impact of regional and local oceanographic processes on community dynamics. The associations from deeper waters harbored more diverse microbial assemblages, and shallow waters, on the other hand, possess higher absolute abundance of microbial cells, suggesting niche specialization across depths. This study underscores the importance of environmental gradients as well as local oceanographic processes in shaping microbial diversity, providing valuable insights into the ecological dynamics of the SB, which are essential for understanding the potential impacts of human activities, such as oil and gas exploration and production, on these critical marine ecosystems.

## INTRODUCTION

Pelagic microbial communities are essential to marine ecosystems, driving processes like nutrient cycling and primary production[1,2]. These communities are pivotal to global biogeochemical cycles, influencing carbon sequestration and nutrient fluxes, and thus sustaining ocean health[3,4]. As marine environments face increasing threats, there’s a pressing global need to characterize marine microbiomes[5,6]. Understanding the connection between microorganisms and local oceanographic processes is crucial for further assessing the impact of human activities[7,8]. Microorganisms can serve as natural biosensors, providing insights into the health of marine ecosystems[6]. The complex dynamics of these microbial assemblages are shaped by a multitude of factors, including physical oceanographic processes, chemical gradients, and biological interactions[9].

Methodological advancements, particularly in molecular techniques such as DNA metabarcoding of the 16S rRNA gene, have revolutionized the study of pelagic microbial communities[10]. These techniques enable researchers to analyze community composition at unprecedented resolutions, revealing intricate biogeographic patterns and ecological interactions[11]. Large-scale oceanographic expeditions like Tara Oceans and Malaspina have provided extensive DNA sequencing data, uncovering the vast microbial diversity present in the epi-, meso-, and bathypelagic realms[12–15]. These studies have identified key environmental drivers, including water masses, temperature, salinity, and nutrients, as crucial factors influencing the geographical and vertical distribution of planktonic prokaryotes[12,15,16].

Recent studies have underscored the significant spatial and temporal variability in microbial communities across oceanic regions[17–22]. Spatial heterogeneity often outweighs temporal variability, as demonstrated by Fortunato et al. [17], where differences across habitats were more pronounced than seasonal fluctuations in bacterioplankton communities. Yet, the vastness and complexity of marine ecosystems, coupled with the challenges of analyzing large-scale molecular datasets, hinder our ability to fully characterize and predict microbial community responses to environmental changes[8,23]. Traditional methods of sample grouping based on predefined factors may fail to capture the true structure and diversity of microbial communities, as they may overlook subtle patterns and interactions influenced by a multitude of environmental variables.

Machine learning offers a powerful solution to address these challenges. By integrating unsupervised methods like self-organizing maps (SOMs) with supervised classification algorithms, we can enhance our ability to predict community responses to environmental changes[24]. These techniques enable the identification of natural groupings within the data, independent of predefined categories, and the detection of subtle patterns and associations within high-dimensional data, offering deeper insights into the environmental factors that drive microbial community structure and making it a significant advantage in the study of marine microbial ecology [25,26].

This study aimed to predict the horizontal and vertical spatial distribution of planktonic microbial communities during two sampling periods in the Santos Basin (SB). We hypothesized that the spatial distribution of microbial community structure is driven by the distinct characteristics of Basin locations, with environmental factors and seasonal variations exerting additional influence on microbial composition and dynamics. Specifically, the objectives were to: (1) model the abundance and composition of planktonic Bacteria and Archaea communities, (2) assess the horizontal and vertical stratification of these communities across the epipelagic to bathypelagic zones, and (3) predict the oceanographic variables driving microbial community composition. To achieve these objectives, seawater samples were collected from multiple stations during two distinct sampling periods within the SB, Brazil’s largest marine sedimentary basin. Covering an area of 350,000 square kilometers, the SB is a critical region for ecosystem services, including genetic resources, fisheries, energy production by oil extraction, and mineral extraction.

## MATERIALS AND METHODS

### Study area

The Santos Basin (SB) covers an area of approximately 350,000 km² along the southeastern-southern Brazilian continental margin, between the parallels 23° and 28°S. The northern boundary is the Cabo Frio High, at Cabo de São Tomé (22.1°S and 41°W), while the southern boundary is the Florianópolis High, near the Pelotas Basin, at Cabo de Santa Marta (28.55°S and 48.47°W)[27,28]. Additionally, the basin extremes are influenced by local and periodic oceanographic processes that support biological productivity. In the northern limit of basin near Cabo Frio High, a coastal upwelling of South Atlantic Central Water (SACW) frequently occurs due the geomorphology, coastline shape and prevalence of northeasterly winds during the summer[29–31]. At the opposite limit, the de la Plata River plume reaches the southern of the basin due to ocean circulation during the austral winter[32,33].

#### Sampling Strategy

Seawater samples were collected at 60 oceanographic stations along the SB during 13 expeditions, encompassing the mainly the winter and spring of 2019 and summer of 2021 (more details in Moreira *et al.,* 2023) onboard the R/V Ocean Stalwart (**Fig. 1**). A total of 366 samples were analyzed, with 174 samples corresponding to the year 2019 and 192 to 2021. Oceanographic stations were distributed in 8 transects perpendicular to the coastline, including the inner, middle and outer shelf, in addition to the continental slope and the oceanic region, reaching up to 2,400 m. Six sampling depths per oceanographic station were predefined according to the water masses nuclei based on the literature, except the DCM (deep chlorophyll maximum), which were defined *in situ* with chlorophyll (RFU) profiles provided by a combined Sea-Bird CTD/Carrousel 911 10l Niskin rosette system, in order to sample the water masses within the epi-, meso- and bathypelagic zones. The depths studied included the surface (SRF) and DCM layers within the epipelagic zone, 250 m and 900 m within the mesopelagic zone (MES), and 1,200 m and 2,300 m within the bathypelagic zone (BAT). Further details on these depths and the associated scientific cruises can be found in Moreira et al. (2023). Seawater (up to 15L) was filtered using a peristaltic pump through 0.22 μm membrane pores (Sterivex, Millipore, MA) for molecular biology. Triplicate samples (1.5 ml) for flow cytometry were stored into cryovials and immediately preserved with Sigma-Aldrich glutaraldehyde (0.1% of final concentration), and flash-freezing in liquid nitrogen. All samples for biological analysis were stored at –80 °C for up to 30 days until analysis. Samples for inorganic nutrients (nitrate, nitrite, phosphate, and silicate) were obtained from the filtered 0.22-μm water, stored at −20 °C, and determined on land by using a flow injection auto-analyzer (Auto-Analyzer 3, Seal *Inc.*). Additionally onboard, seawater (1-5L) from the SRF and DCM was filtered using a vacuum pump through 0.7 μm fiberglass membrane (Whatman GF/F) and stored in liquid nitrogen for chlorophyll-a (CHL-a) measurement. On land, fiberglass membranes were extracted with 90% acetone and determined by using Turner 10AU fluorometer[34].

**Figure 1.**
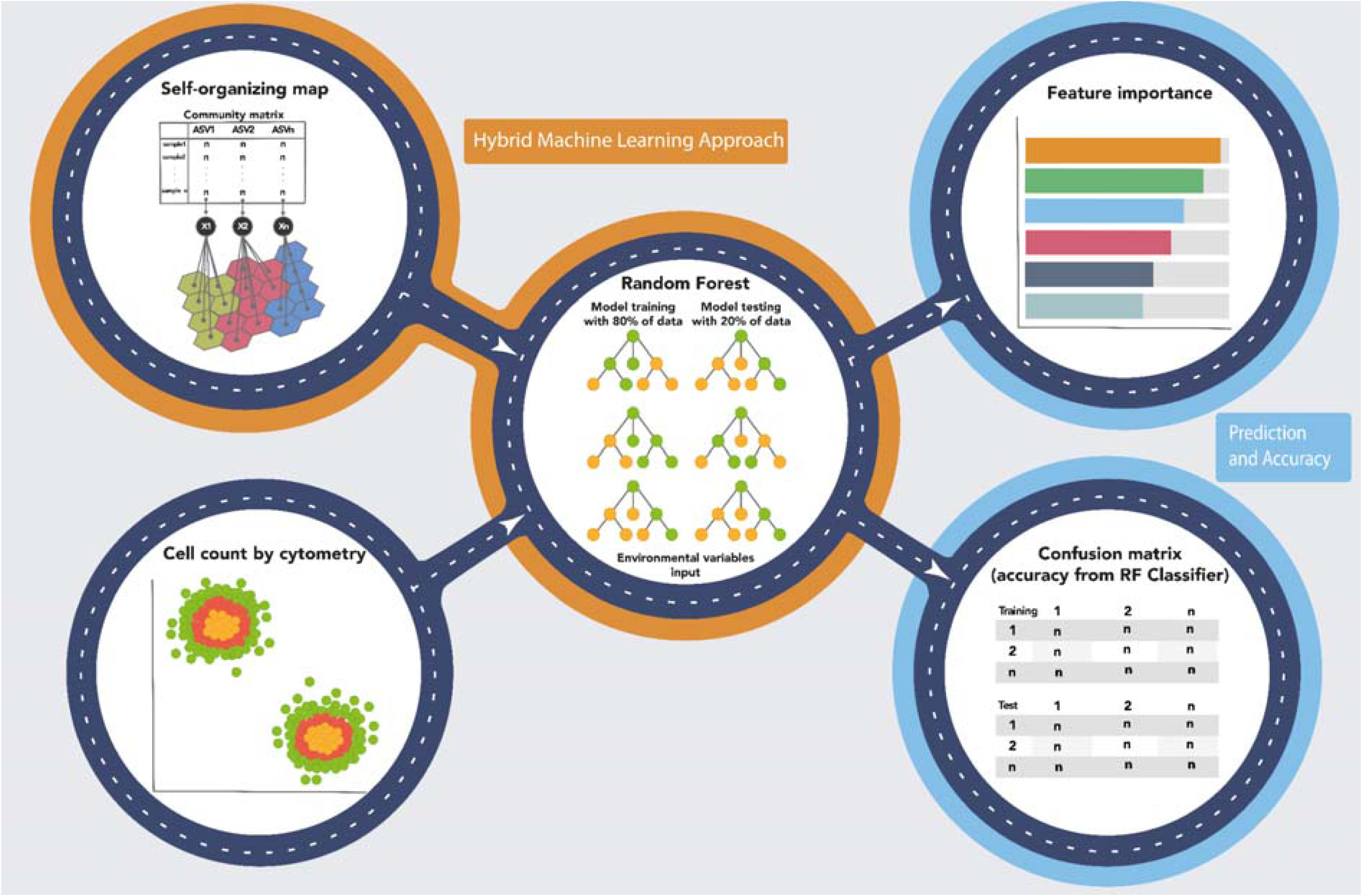
Hybrid machine learning approach integrating self-organizing maps (SOM) and random forest (RF) classification for microbial community analysis. The workflow consists of multiple steps: (i) a self-organizing map (SOM) (top-left) is used to cluster microbial communities based on an input matrix representing the relative abundance and occurrence of taxa across samples; (ii) a random forest model (center) is trained using 80% of the data and tested on 20%, with environmental variables as input for classification; (iii) the feature importance analysis (top-right) highlights the most influential environmental variables in the classification model; (iv) a confusion matrix (bottom-right) evaluates the RF model’s performance, comparing predicted versus actual classifications for training and test datasets; and (v) cytometric cell counts (bottom-left) provide additional microbial community data, visualized through a scatter plot where colors represent different microbial clusters.

#### Flow cytometry analysis

Cell abundance of heterotrophic prokaryotes, *Synechococcus* spp., *Prochlorococcus* spp., picoeukaryotes (≤2 μm), and nanoeukaryotes (2–5 μm) were determined using an Attune NxT flow cytometer (Thermo Fisher Scientific), following the methods of Marie et al.[35]. The flow cytometer was equipped with a syringe pump for quantitative volume sampling and two flat-top lasers (blue at 488 nm and red at 640 nm). *Prochlorococcus* spp.*, Synechococcus* spp., pico and nano-eukaryotes were quantified based on their autofluorescence at red (BL3 - intracellular chlorophyll concentration, 656 nm) and orange (BL2 - intracellular phycoerythrin concentration, 530 ± 30 nm) wavelengths simultaneously. To reduce noise, a threshold of 500 was applied to BL3. After autotroph analysis, SYBR Green I (Invitrogen Life Technologies, USA) was added, and samples were dark-incubated at room temperature for 15 minutes. Flow cytometry acquisition for heterotrophic prokaryotes was triggered on BL1 (530 ± 30 nm) with a threshold value of 500 and 90° side scatter. To estimate the total abundance of prokaryotic cells, the counts of heterotrophic bacteria, *Synechococcus* spp., and *Prochlorococcus* spp. were summed. This combined total represents the overall prokaryotic community, encompassing both autotrophic (*Synechococcus* and *Prochlorococcus*) and heterotrophic components.

#### DNA Extraction, 16S rRNA Gene Amplification, and Illumina Miseq Sequencing

DNA extraction was performed by using the DNeasy PowerWater (Qiagen, EUA) according to the manufacturer’s instructions. Negative extraction controls (no sample added) were used to ensure the extraction quality. DNA integrity was determined after electrophoresis in 1% (v/v) agarose gel prepared with TAE 1X (Tris 0.04M, glacial acetic acid 1M, EDTA 50mM, pH 8), and staining with Sybr Green safe (Thermo Fisher Scientific, São Paulo, Brazil). DNA concentration was determined using the Qubit dsDNA HS assay kit (Thermo Fisher Scientific, São Paulo, Brazil), according to the manufacturer’s instructions. Illumina DNA libraries and sequencing were performed on a MiSeq platform in a paired-end read run (2 × 300bp) following the manufacturer’s guidelines. The V3-V4 region of the 16S rRNA gene was amplified with the primer set 515F (5’– GTGCCAGCMGCCGCGGTAA-3’) and 926R (5’– CCGYCAATTYMTTTRAGTTT-3’)[36]. Negative DNA extraction controls (no sample added) were used for PCR amplification and Illumina sequencing to check for the presence of possible contamination during DNA extraction procedure.

#### Data Processing and Diversity Analysis

For sequencing data analysis, the demultiplexed sequences were processed using the software package Quantitative Insights Into Microbial Ecology (QIIME 2) version 2019.4[37]. Sequences were denoised using DADA2[38] with the following parameters: trim left-f = 24, trim left-r = 25, trunc-len-f = 260, trunc-len-r = 220. The taxonomy was assigned to the representative sequences of ASVs using a Naive Bayes classifier pre-trained on SILVA release 138 clustered at 99% identity. Alpha-diversity metric (Shannon diversity) was calculated using repeated rarefaction (500 iterations) to account for variations in sequencing depth across samples[39]. We performed repeated rarefaction at a sequencing depth of 6,700 reads, the minimum threshold required to include all samples. The diversity indices were compared between groups using Wilcoxon’s rank test for paired samples (Holm’s method adjustment).

#### Hybrid Machine Learning Pipeline

The sequencing dataset at the family level was implemented in the hybrid approach (**Fig. 1**) that integrated both unsupervised and supervised learning methods (detailed below) through the application of iMESc[24]. The analysis was conducted at the family level to reduce data dimensionality and computational complexity.

##### Unsupervised Learning

For the unsupervised analysis, we utilized self-organizing maps (SOMs), a neural network technique[40], to group similar samples into neurons, known as Best-Matching Units (BMUs). In this study, a multi-layered SOM approach was employed. Given the high diversity of family (580 families) identified from the annotated reads, we transformed the family-level data using the square-root method. The first SOM layer emphasized microbial family-level data with a weight of 0.95, based on the Euclidean similarity index. The second layer integrated spatial coordinates and depth information (scaled and centered), weighted at 0.05, also using Euclidean distance. The number of SOM nodes was determined using the default set of iMESc, which follows Vesanto’s[41] guidelines. Each neuron in the SOM map represents a weighted list of microbial families, referred to as a codebook. The contribution of each microbial family was assessed by summing their absolute weights, with the top 10 families identified. The codebooks were then subjected to hierarchical clustering (Ward’s method with squared differences, “Ward2”) to group similar neurons and their corresponding best-matching units (BMUs). The number of clusters was determined by examining the relationship between cluster number and within-cluster sum of squares (WSS). As suggested by Vieira et al.[24], these neuron groups, termed taxonomic associations (simply Associations), represent the microbial communities. Shannon and Chao1 alpha diversity indices were calculated for each Association compared across by means of Wilcoxon’s rank test (Holm’s method adjustment). To identify the indicator families for each taxonomic association, the “indicspecies” package[42] was used, applying the “IndVal.g” method. The top five indicator families for each association were retained by selecting those with the highest indicator values.

#### Supervised Learning

In the supervised phase, each taxonomic association was treated as a response variable to identify the optimal set of environmental features for modeling and predicting microbial community associations. Random Forest (RF) classification algorithm was employed, with data divided into training (80%) and test (20%), ensuring balanced representation across the associations. The RF model used 500 trees and a 5-fold cross-validation method repeated 5 times, with hyperparameter tuning performed across a grid search with five candidate values (2, 5, 9, 12 and 16 variables randomly sampled for model evaluation). The model performance was evaluated through accuracy, confusion matrix analysis, and feature importance assessment, highlighting the environmental variables that most significantly influenced the SB microbiome associations. To establish a baseline for model performance, we implemented a stratified classification approach, generating predictions based on the class distribution in the training data. This approach provided a minimal benchmark for evaluating the efficacy of supervised learning models. To assess the significance of the improvement achieved by the RF model over the baseline, we used a bootstrap-based confidence interval for accuracy differences and the McNemar test for patterns of error, ensuring robust statistical validation.

The influence score, reflecting both positive and negative impacts, was calculated by multiplying the variable importance by the correlation coefficient between the environmental data and the class probabilities.

A RF regression model was implemented to analyze the absolute abundance of *Synechococcus*, *Prochlorococcus*, heterotrophic bacteria, and total prokaryotic cells, as measured by flow cytometry, with the aim of identifying the optimal set of environmental features for modeling and predicting microbial community associations. The model setup, including the training and test data partitioning and model tuning followed the same approach described earlier for RF classification. The response variables were log-transformed using base-10 logarithms to normalize their distributions and mitigate the influence of outliers. A naïve regressor, generating random predictions from a normal distribution parameterized by the mean and standard deviation of the training data, was employed as a baseline for comparison. The improvement of the RF model over the baseline was assessed through a bootstrap-based approach to compare differences in Root Mean Square Error (RMSE), a metric that quantifies the average magnitude of prediction errors in regression models. A 95% confidence interval (CI) was calculated to determine the statistical significance of the RMSE improvement.

### Data and code availability

Raw sequence data generated for this are publicly available in the National Centre for Biotechnology Information (NCBI) database under the BioProject PRJNA1172717.

For verifiability of this work, the processed data table (matrix of microbial family abundances) used as input for all analyses are available at Sup. Data S1, as well as the metadata on Sup. Data S2. The iMESc Savepoints used in the analysis are available <u>here</u> and can be used to reproduce the results and figures. The codes for reproducing the alpha diversity results and figures, as well as the machine learning pipeline, are available at [https://github.com/DaniloCVieira/SantosBasin-Microbiome-ML].

## RESULTS

### General Features of Microbial Composition Across Santos Basin

The taxonomic profiling, at phylum level, across all samples collected during 2019 and 2021 expeditions showed an apparent enrichment of *Bacteroidota* and *Planctomycetota* in 2019 compared to 2021 (**Fig. 2B**). On the other hand, 2021 showed an enrichment of ASVs assigned to *Marinimicrobia* (SAR406 clade). In general, the *Proteobacteria*, *Cyanobacteria*, and *Bacteroidota* phyla were substantially abundant in the SRF and DCM. Conversely, the MES and BAT were enriched with ASVs assigned to the *Crenarchaeota* and *Planctomycetota* phyla (**Fig. 2B**). To further explore the dominant communities, we selected the 30 most abundant (relative abundance) families for a detailed examination of their distribution (**Fig. S1 and S2**). Among them, we identified microbial families with high abundance at the SB, such as Clade I (SAR11_clade), *Cyanobiaceae*, and *Actinomarinaceae* which had higher abundance mainly on SRF waters (**Fig. S1 and S2**), while *Nitrosopumilaceae* was more abundant in the MES and BAT zones (**Fig. S1 and S2**).

**Figure 2.**
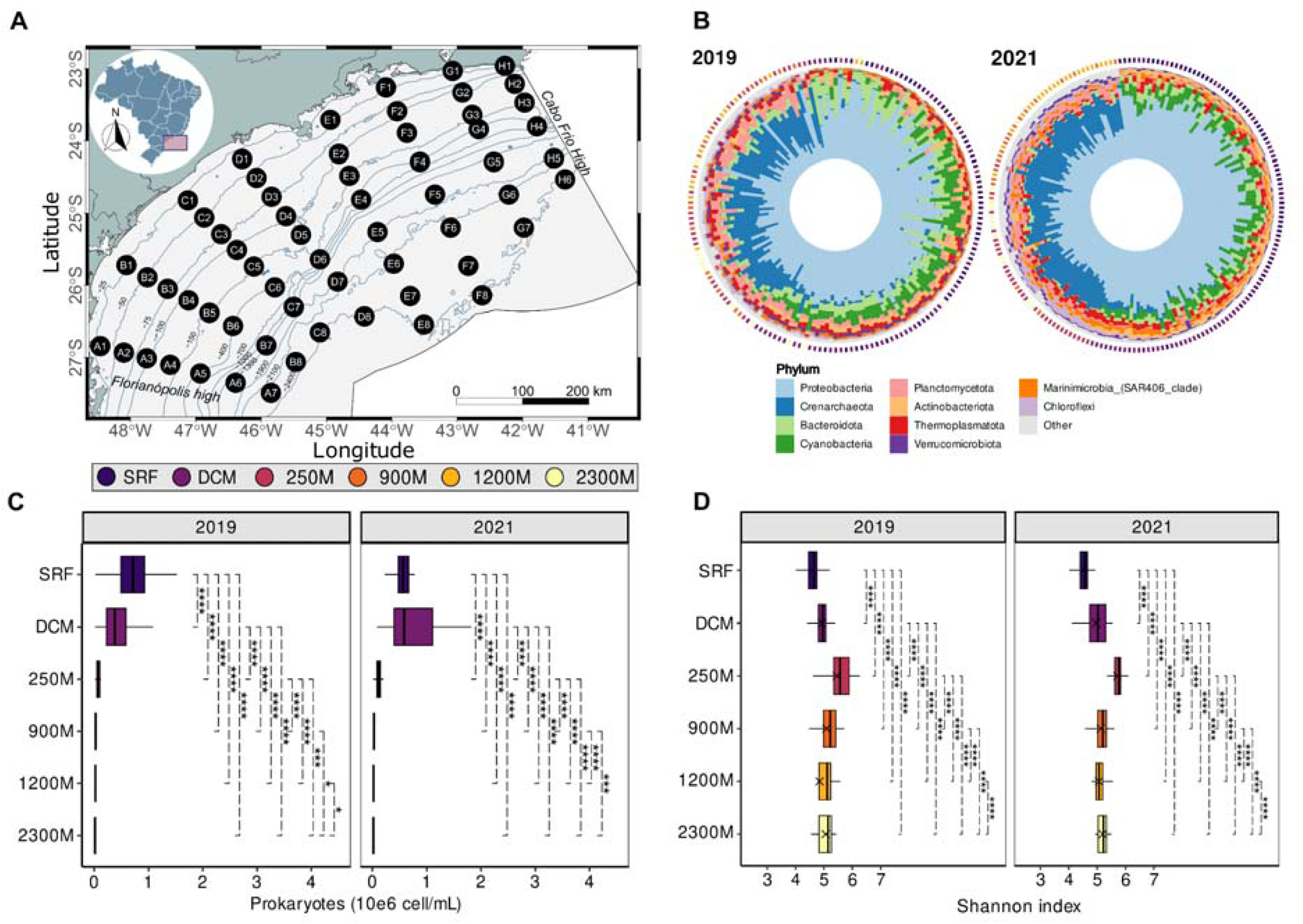
Spatiotemporal variations of the microbial community structure and diversity in the Santos Basin. (A) Sampling map of the Santos Basin, showing oceanographic station locations (numbers) and transects (letters). . (B) Taxonomic distribution of microbial communities at the Phylum level in the Santos Basin, comparing data from 2019 (left) and 2021 (right), sorted by pelagic zones. (C) Prokaryotic cell counts measured by flow cytometry. Higher cell count values indicate greater absolute abundance level. (D) Shannon index values representing the diversity of microbial communities across different pelagic zones and years, measured by amplicon sequencing data. Higher Shannon index values indicate greater diversity. Significance levels (p-adjusted) are indicated by: ns: p > 0.05; *: p ≤ 0.05; **: p ≤ 0.01; ***: p ≤ 0.001; ****: p ≤ 0.0001. Error bars represent the standard deviation of the mean. Statistical comparisons were performed using Wilcoxon’s rank test.

In both years, the cell count by cytometry showed there was a clear enrichment in absolute abundance levels of total prokaryotic cells in the epipelagic zone (SRF and DCM samples; **Fig. 2C**). In 2019, SRF samples showed a significantly higher abundance compared to DCM samples (Wilcoxon, p-adjusted ≤ 0.0001). However, in 2021, there was an increase in absolute abundance for DCM samples, with no significant difference compared to SRF samples (**Fig. 2C**), suggesting a shift in abundance depending on the sampling period.

Interestingly, the alpha-diversity measurement (Shannon Index) revealed different patterns compared to absolute abundance. There was a clear increase in Shannon Index values at 250m depth for both years (**Fig. 2D**), in contrast to other sampling depths. Additionally, greater depths exhibited higher Shannon Index values compared to SRF and DCM samples, indicating an increase in diversity with increasing depth.

### Predicting Spatial Profile of Santos Basin Microbiome by Machine Learning

The SOM analysis stabilized with a learning rate of approximately 0.0017 for ASV data assigned to family level and 0.03 for geographical data. The SOM network accounted for 72.9% of the data variance, with a mean topographic error of 0.05. Best-Matching Units (BMUs) exhibited a pattern according to depth (**Fig. 3A**), highlighting depth-stratified microbial associations. The top 10 families with the highest absolute weights in the SOM codebook (**Fig. 3B**) included Clade I (SAR11_clade), *Nitrosopumilaceae*, Clade II (SAR11_clade), SAR86 clade, *Flavobacteriaceae*, Marine Group II (*Thermoplasmata*), *Cyanobiaceae*, *Marinimicrobia* (SAR406), NS9 Marine Group (*Flavobacteriales*), and AEGEAN-169 Marine Group (*Rhodospirillales*) (Figure 2C-L). *Cyanobiaceae* and AEGEAN-169 Marine Group (*Rhodospirillales*) were primarily distributed in the photic surface waters (SRF and DCM), consistent with their epipelagic zone dominance. *Flavobacteriaceae* and NS9 Marine Group (*Flavobacteriales*) were mainly concentrated in the twilight zone (250M), however, with substantial abundance observed in the photic DCM. Clade I (SAR11_clade), Clade II (SAR11_clade), SAR86 clade, and Marine Group II (*Thermoplasmata*) exhibited broad distributions across multiple depths, with notable presence in the photic and twilight waters (SRF, DCM and 250M). *Nitrosopumilaceae* and *Marinimicrobia* (SAR406) were predominantly found in the dark and deeper waters, with *Marinimicrobia* concentrated in mesopelagic samples (250M and 900M), while *Nitrosopumilaceae* showing higher relative abundance in the bathypelagic samples (900M and 2300M) .

**Figure 3.**
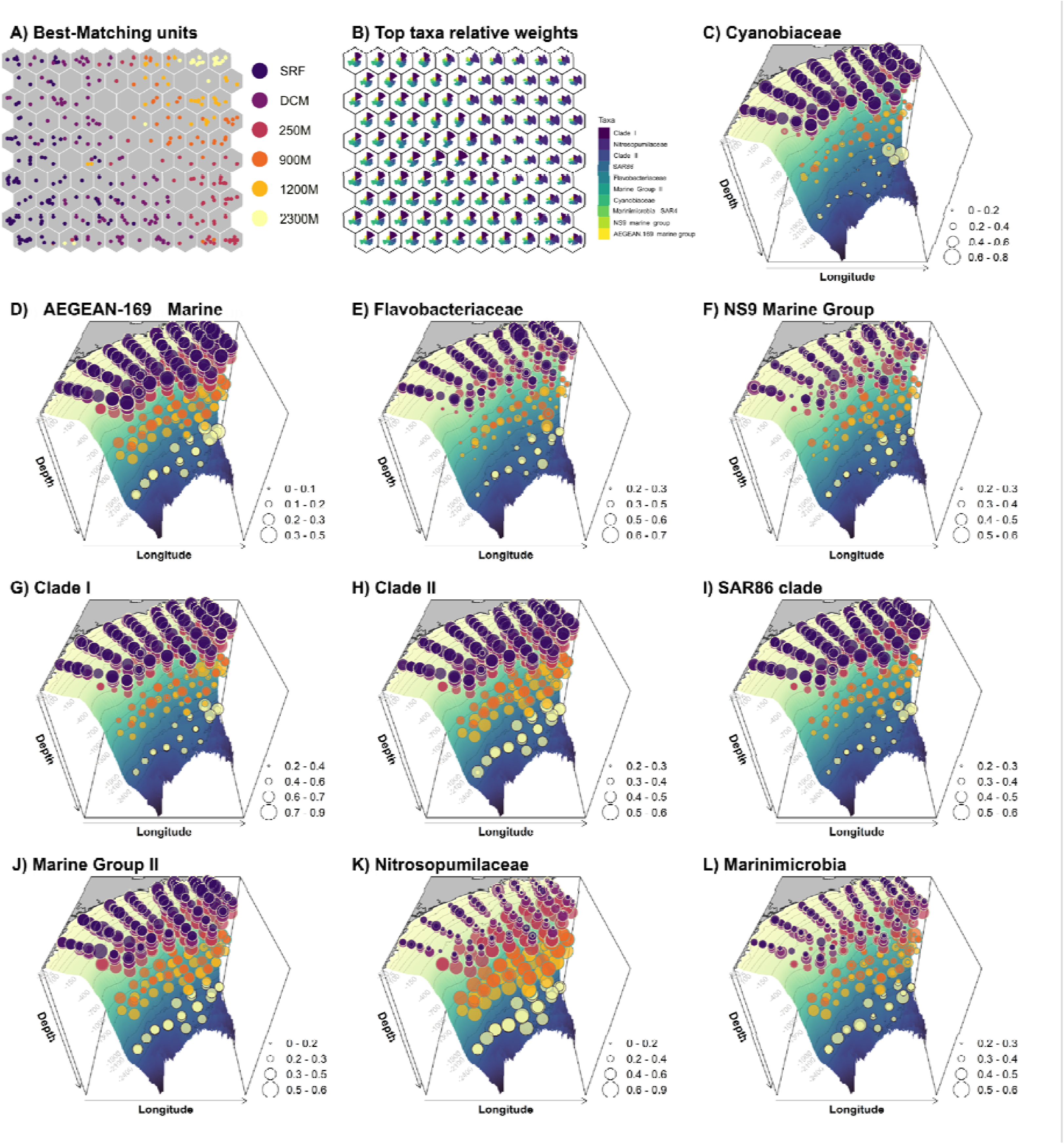
Spatial distribution of top marine taxa in the Santos Basin revealed by Self-Organizing Maps (SOM) and measurement of absolute weights. **(A)** Best-Matching Units (BMUs): Each hexagon represents a neuron in the SOM, and only hexagons containing samples are considered Best-Matching Units (BMUs). BMUs are colored by depth. **(B)** Top taxa with the highest absolute weights: The size of the pie charts reflects relative absolute weights in the SOM codebook. **(C–L)** Three-dimensional distribution of the top taxa with high absolute weight in the Santos Basin. The square root (sqrt) of the relative abundance of each family is represented by the circle diameter. The gradient colors, transitioning from yellowish to bluish, indicate the depths within the Santos Basin, where yellow represents shallow waters and deep blue indicates deeper ocean regions.

Hierarchical clustering from SOM’s algorithm identified five taxonomic associations, representing different microbial assemblages, (A1-A5; **Fig. 4A**) with distinct annual, spatial and vertical distribution patterns across the SB. The yearly photic spatial distribution (Associations 1, 2 and 3, **Fig. 4B, D**) were observed in the SB epipelagic zone: (i) The southern continental region in 2019 was represented by southern coastal SRF and DCM samples and, was represented by only southern coastal DCM samples in 2021 (A1, purple association); (ii) Coastal resurgence in 2021 was represented by two northern coastal SRF samples (Cabo Frio and Guanabara Bay) and coastal DCM samples (A1, purple association); (iii) SRF samples (A2, blue association) was prevalent in the northern portion of SB in 2019, and was widespread across SRF samples in 2021; (iv) Prevalent epipelagic oceanic samples (A3, light green association) appeared in southern SRF samples in 2019 and exhibited similar patterns in DCM samples from 2019 and 2021. The twilight and dark ocean distribution (vertical patterns, A4 and A5, **Fig. 4B, D**) were not influence by yearly local oceanographic processes: (i) twilight transition zone was prevalent in samples from 250m (A4, light orange association); (ii) dark ocean was abundant in deeper samples (< 900m, A5, dark red).

**Figure 4.**
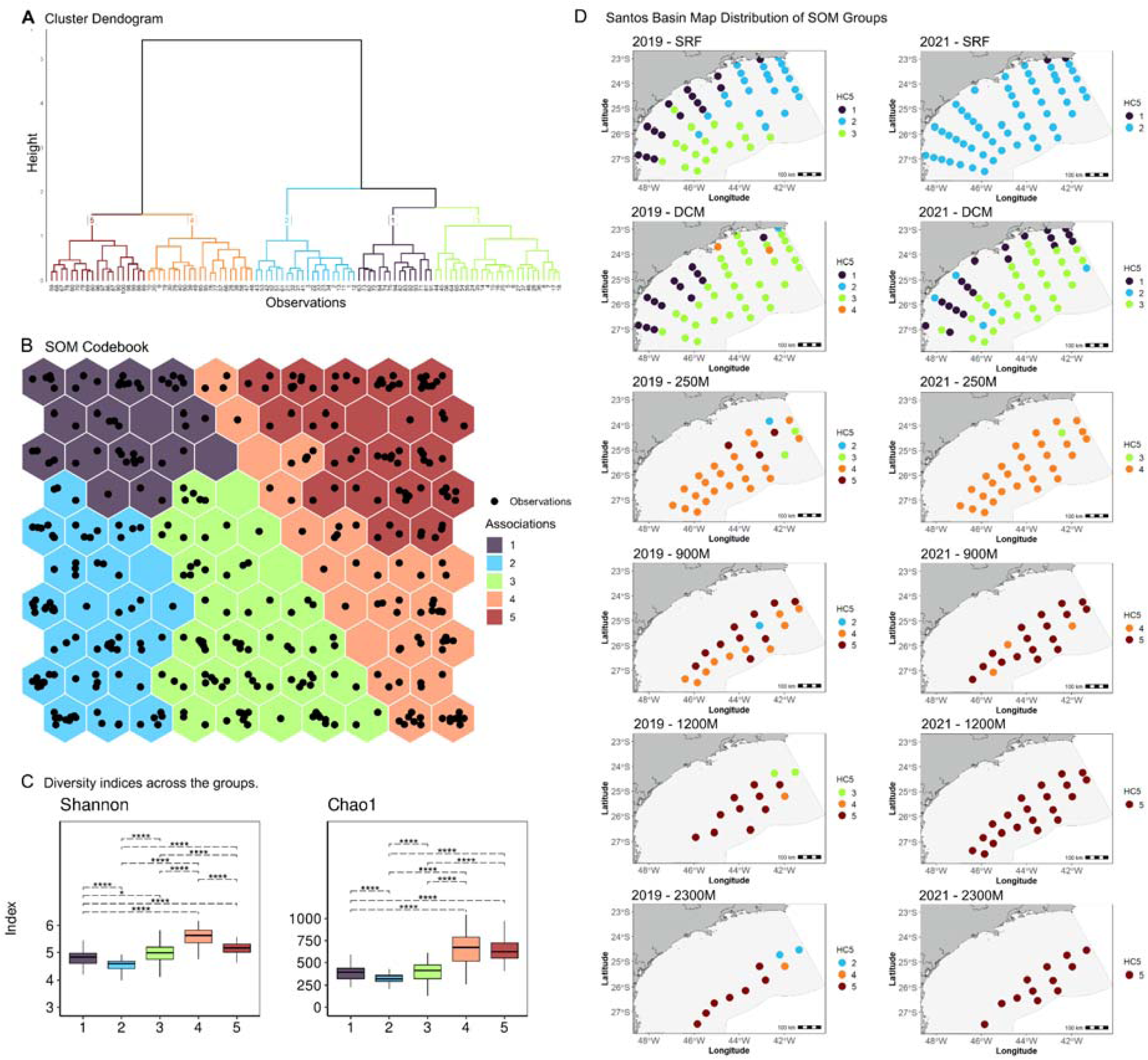
A). Cluster dendrogram showing the division of observations into five distinct taxonomic associations (Groups 1 to 5). B) SOM codebook. Neurons are colored by their assigned group based on hierarchical clustering. (C) Boxplots of Shannon and Chao1 diversity indices for each of the five groups. (D) Santos Basin map distribution of SOM groups for 2019 and 2021 across depth zones. Higher Shannon index values indicate greater diversity whilst higher Chao1 values indicate higher richness. Significance levels (p-adjusted) are indicated by: ns: p > 0.05; *: p ≤ 0.05; **: p ≤ 0.01; ***: p ≤ 0.001; ****: p ≤ 0.0001. Error bars represent the standard deviation of the mean. Statistical comparisons were performed using Wilcoxon’s rank test.

The **IndVal.g** results identified key indicator taxa, with distinct distribution patterns across the SB (**Figs. S4-S8**). SOM association 1 indicators included *Methylophilaceae*, NS11-12 Marine Group (*Sphingobacteriales*), OM182 Clade, *Pedosphaeraceae*, and PeM15 (*Actinobacteria*) (**Fig. S4**). *Methylophilaceae* and PeM15 showed higher relative abundance on coastal samples in contrast to oceanic samples, for both years (**Fig. S4**). On the other hand, *Pedosphaeraceae* was widespread through southern coastal surface and DCM water in 2019, and more restricted to northern coastal DCM samples in 2021. The families assessed by higher IndVal.g values on SOM’s Association 2 (Clade III - SAR11_clade), *Litoricolaceae*, *Microcystaceae,* MWHUniP1 aquatic group and PS1 Clade) showed a similar trend: they were evenly distributed across the entire surface water area of the SB in 2019 but concentrated in the northern region in 2021 (**Fig. S5**). As for the families assigned to Association 3 (Clade IV - SAR11_clade, *Cyclobacteriaceae*, *Microcystaceae*, DEV007, and OCS 116 Clade), although they exhibited higher relative abundance at the DCM depth in both years, notably, all five families were present across all SOM associations and depths (**Fig. S6**), indicating a broad adaptability throughout the water column. The indicator families assigned to SOM Association 4 were *Dehalococcoidia* (Class), *Planctomycetes* (Class), Marine Benthic Group A (*Nitrososphaeria* Class), *Nitrospiraceae* and SAR11 Clade. These families exhibited similar patterns of occurrence and relative abundance across the years analyzed, with a predominance in mesopelagic waters (mainly in the 250 m than 900 m samples). Except for *Dehalococcoidia* with higher abundance in 2021 and the *Nitrospiraceae* family with more dominance in the 250 m samples (**Fig. S7**). Conversely, the taxa assigned to Association 5 (Arctict95B-14, *Moritellaceae*, *Woesearchaeales*, PAUC26f, and SM23-30) were enriched in samples from depths ranging from 900 to 2300 m but also appeared with lower occurrence and relative abundance in the 250 m samples (**Fig. S8**).

### Environmental Properties on Microbial Communities

#### Predictive performance of RF model and accuracy

The Random Forest (RF) model achieved an accuracy of 85.10% on the training dataset with cross validation and 86.11% using the final model on the test dataset (**Fig. 5A**). In comparison, the stratified baseline, which generated predictions based on the class distribution in the training dataset, achieved an accuracy of 13.89% on the test dataset. The RF model significantly outperformed the baseline, as confirmed by McNemar’s test (χ² = 46.446, p < 0.001), indicating a substantial improvement in predictive performance.

**Figure 5.**
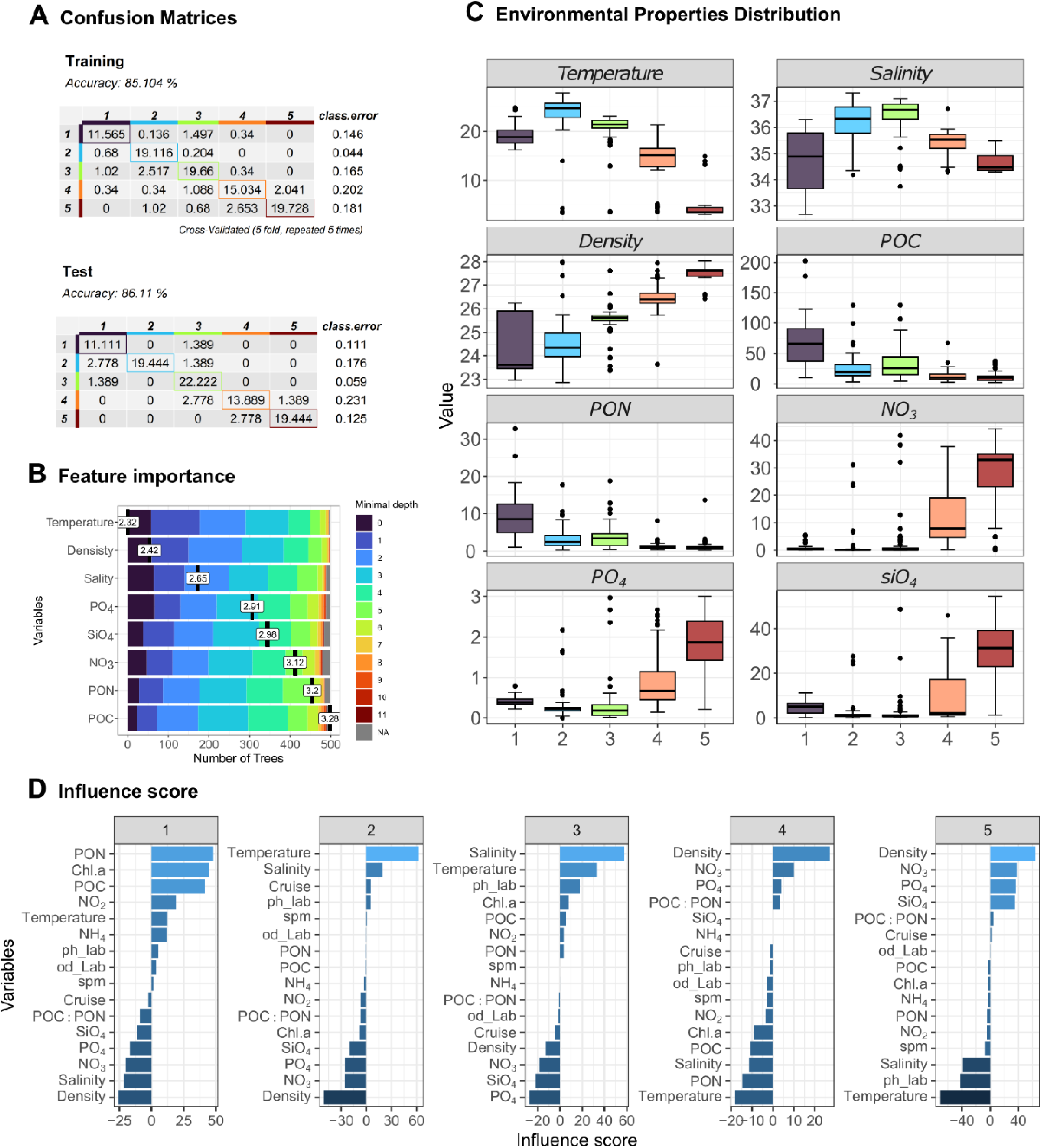
Model Performance, Variable Importance, and Environmental Influence. A: Classification accuracy and errors for training and test datasets, with actual vs. predicted classes. B: Importance of environmental variables in the Random Forest model, colored by minimal tree depth. C: Boxplots showing the distribution of key environmental properties across the five classes. D: Influence scores for each variable across the five classes, reflecting the direction and magnitude of their impact on class predictions.

Associations 1 and 2 had the lowest error rates (14.6% and 4.0%, respectively) in the training data, while Associations 4 and 5 had the highest error rates (20.2% and 18.1%, respectively). Association 3 showed 16.5% of error rate on training data. In the test data, on the other hand, Association 3 had the lowest misclassification rate at 5.0%, followed by Association 1 at 11.1%, Association 5 at 12.5%, and Association 2 at 17.6%. Association 4 presented the highest misclassification rate at 23.1% on test data (**Fig. 5A**).

#### Estimating the importance of environmental variables on microbial communities

Among the 16 environmental variables, eight were selected as significant (p-value < 0.05) (**Fig. 5B**). Temperature emerged as the most important feature, followed by density, salinity, PO_4_, SiO_4_, NO_3_, PON, and POC (**Fig. 5B**). Higher temperatures were observed in association 2 compared to the other associations (**Fig. 5C**), with a slight decrease in association 3 and association 1. Lower temperature was noted in Association 5. Salinity followed a similar trend to temperature (**Fig. 5C**); however, association 1 exhibited considerable variability in salinity measurements compared to the other associations. In fact, association 1 displayed distinct patterns in measurements of other environmental properties, including POC, PON, PO_4_, and SiO_4_ (**Fig. 5C**).

Further analysis using influence scores highlighted the differential impact of these environmental variables on microbial community dynamics across the five SOM associations (**Fig. 5D**). For association 1, PON and chlorophyll-a were the primary positively-associated environmental factors, while density and salinity exerted the most significant negative influences. In contrast, association 2 displayed a positive influence from temperature and salinity, with density and NO_3_ showing the highest negative influence scores. Association 3 was positively associated with salinity and temperature, while PO_4_ and SiO_4_ were the most negatively-associated environmental factors. As expected for association 4, the pattern shifted, with density, NO_3_, and PO_4_ emerging as the most positively-associated factors, while temperature and PON had the higher negative influence. Similarly, association 5 was positively influenced by density and NO_3_, while temperature and pH (lab) were the most prominent negative factors.

#### Environmental Properties and Temporal Variation Shape the Absolute Abundance of Planktonic Microorganisms

For the cytometry cell-count data (absolute abundance), the R² values based on training data ranged from 0.61 to 0.71 across different microbial taxa, while test R² values ranged from 0.619 to 0.691. The Random Forest (RF) model consistently outperformed the naïve baseline, as demonstrated by lower RMSE values and statistically significant improvements in predictive performance (p < 0.05, based on bootstrap confidence intervals for RMSE differences). For Heterotrophic bacteria, the training R² was 0.61 and the test R² was 0.662, with an RMSE of 0.400 compared to 0.689 for the naïve baseline. The improvement was statistically significant (95% CI for RMSE difference: −0.429 to −0.169). The RMSE metric indicates better model performance with lower values. The interval (−0.429, −0.169) reflects that the RF model’s RMSE is consistently lower than the baseline, as negative values denote superior RF performance. The model identified water density, temperature, and NO_3_ as the most influential features that explains the absolute abundance of planktonic microorganisms (**Fig. S9**). These variables were critical in determining the abundance patterns observed, with temperature showing the highest positive influence scores and density with the highest negative influence (**Fig. S10**). By contrast, PO_4_ and density were the most significant negatively-associated factors (**Fig. S10**).

For *Prochlorococcus*, the model achieved a training R² of 0.64 and a test R² of 0.665, with an RMSE of 0.623 compared to 1.060 for the naïve baseline. The improvement was significant (95% CI for RMSE difference: −0.603 to −0.271). The top predictors included pH (lab), density, and temperature emerging as the top predictors (**Fig. S9**). Temperature, again, played a pivotal role as a positive driver of *Prochlorococcus* abundance, together with the pH and the levels of Chlorophyll-a, while density and PO_4_ exhibited strong negative influence scores (**Fig. S10**), implying that these factors might limit the growth of this group under certain conditions. Similarly, for *Synechococcus*, the RF model (training R² = 0.65, test R² = 0.691) highlighted density and salinity as key determinants (**Fig. S9**), with PON, salinity, pH and temperature exerting the highest positive influence (**Fig. S10**).

The model’s performance was highest for the total prokaryotic community, with a training R² of 0.71 and a test R² of 0.619. The RF model achieved an RMSE of 0.445, compared to a naïve RMSE of 0.720, with a significant difference confirmed by a 95% confidence interval [−0.417, −0.151]. Temperature, density and NO_3_ identified as the most important variables (**Fig. S9**), which is aligned with the findings within the RF model for the amplicon sequencing data. Temperature, PON, chlorophyll-a, salinity and pH showed the most substantial positive influence on prokaryotic absolute abundance, while density, PO_4_, and NO_3_ were associated with negative influence scores (**Fig. S10**).

## DISCUSSION

The microbial communities of the Santos Basin (SB) are shaped by a dynamic interplay of abiotic factors, local oceanographic processes, and temporal variation, with depth-driven stratification emerging as the dominant organizational principle. Our integration of self-organizing maps (SOMs) and machine learning models revealed that temperature, salinity, and nutrient gradients are the primary drivers of microbial community structure, with locally traits of spatial-temporal patterns exerting additional influences on these gradients. While temporal shifts, such as the increased relative abundance of *Marinimicrobia* (SAR406) in 2021 compared to *Bacteroidota*-enriched 2019 communities (**Fig. 2B**), suggest interannual variability, these changes were tightly coupled to fluctuations in environmental conditions. For instance, the 2021 rise in *Marinimicrobia*, a group associated with oligotrophic, mesopelagic waters and oxygen minimum zone[43,44], coincided with elevated phosphate (POU) and nitrate (NOU) levels (**Fig. 5B–C**), likely driven by intensified Cabo Frio upwelling bringing nutrient-rich South Atlantic Central Water (SACW) to the photic zone[45,46]. Similarly, the dominance of *Methylophilaceae* and *Pedosphaeraceae* in coastal Association 1 (**Figs. S4–S5**) aligned with lower salinity and higher particulate organic nitrogen (PON), reflecting freshwater influences from the La Plata plume[33,47]. These findings challenge earlier studies in marine systems where seasonal cycles dominated the explanation on microbial turnover[48–53], instead highlighting how regional oceanographic features mediate temporal effects through abiotic restructuring.

The vertical partitioning of SB’s microbiome further underscores abiotic control. Epipelagic zones were dominated by taxa such as *Cyanobiaceae* and *SAR11 Clade I*, whose photoheterotrophic strategies[54,55] align with high light availability and warmer temperatures (**Fig. 3C–L**). In contrast, mesopelagic and bathypelagic communities, though more stable temporally, exhibited distinct groups that reflects functional adaptations: *Nitrosopumilaceae* and *Marinimicrobia* in twilight zones (250–900 m) correlated with ammonia oxidation and sulfur cycling[44,56], while bathypelagic *Woesearchaeales* and *PAUC26f* (**Fig. S8**) suggest anaerobic metabolisms fueled by particulate organic matter sinking[57–60]. The higher Shannon diversity in deeper layers (**Fig. 2D**) with diversity peaks at mid-depths which may be driven by water masses traits including the temperature, density, NO_3_ PO_4_ and SiO_4_(**Fig. 5C**). This depth-dependent stability implies that deep-sea communities may buffer environmental change more effectively than surface layers—a critical insight for conservation in a basin facing escalating deep-sea oil exploration[61].

Our machine learning framework resolved hidden spatial-temporal interactions that traditional methods may overlook. For example, SOMs detected a northward expansion of Association 2 in 2021 (**Fig. 4B**), characterized by *SAR11 Clade III* and *Litoricolaceae (syn. Litorivicinaceae)*, which RF models tied to rising temperatures and salinity (**Fig. 5B–D**). These taxa, typically associated with oligotrophic open oceans[54,62], proliferated in SB’s northern SB as SACW encroached, displacing coastal taxa like *PeM15*. Such fine-scale redistributions, missed by conventional beta-diversity analyses, demonstrate how unsupervised learning can disentangle overlapping environmental signals. Similarly, RF’s identification of density and chlorophyll-*a* as key predictors of *Prochlorococcus* abundance (**Fig. S9–S10**) aligns with its known niche in stratified, nutrient-poor waters (*Flombaum et al., 2013*), yet our models uniquely linked these factors to interannual shifts in cell counts (**Fig. 2C**), providing a predictive framework for microbial responses to eutrophication or warming.

However, our study has limitations. While we identified PON and POC as critical drivers of heterotrophic bacteria (**Fig. S6**), for instance, trace metals like iron — key to nitrogen fixation in SB and likely sourced from the La Plata plume[63] — were not analyzed. Future work should integrate multi-omics and geochemical profiling to resolve resource competition and use long-term data collecting for monitoring the community dynamics. Nevertheless, our findings establish that abiotic factors, modulated by oceanographic processes, are the ultimate drivers of microbial structure in SB, with temporal variation acting indirectly through physicochemical restructuring. This mechanistic understanding is vital for forecasting impacts of climate change and industrial activity in this ecologically and economically pivotal region.

In summary, this study highlights that the microbial biogeography of the Santos Basin is primarily shaped by abiotic factors—temperature, salinity, and nutrient gradients—modulated by dynamic oceanographic processes such as the Cabo Frio upwelling and La Plata plume intrusions. While temporal shifts in community composition occur, they are not strictly seasonal but rather emerge from physicochemical restructuring driven by these regional phenomena. Our hybrid machine learning approach uncovered nuanced spatiotemporal interactions, such as the northward expansion of oligotrophic taxa under upwelling influence and depth-dependent stability. These findings challenge conventional paradigms that prioritize seasonal cycles over oceanographic mediation, emphasizing the critical role of vertical stratification and regional hydrography in microbial ecology. Beyond advancing our understanding of microbial community dynamics, our results have implications for predicting microbial resilience to climate change and increasing anthropogenic pressures, such as deep-sea oil exploration. Despite existing knowledge gaps, our framework establishes a foundation for future multi-omics studies aimed at disentangling resource competition and metabolic strategies. Ultimately, this mechanistic perspective positions the Santos Basin as a model system for forecasting microbial responses to environmental perturbations in a rapidly changing ocean.

## Conflict of Interest

The authors declare no conflict of interest.

### Availability of data and material

The sequencing reads generated for this study can be found in the National Centre for Biotechnology Information (NCBI) database under BioProject PRJNA1172717.

## Acknowledgments

We are grateful to Petrobras for the project planning, coordination, execution, and financing of the Santos Project (PCR-BS) and his working package “Caracterização química e biológica do sistema pelágico da Bacia de Santos”. To all researchers and their institutions for being part of this project, nominally: UFRJ, UFF, PUC-Rio, UERJ, FIRJAN/SENAI, SALT, INPE, USP, UNIFESP, UFPR, and others not part of the cruises but equally important UNESP, IP-SP, FURG and Socioambiental. To Foundation in support of the University of São Paulo (FUSP) for administrative management covering equipment purchase and maintenance, scholarship, travel, and others. To the vast crew of RV Ocean Stalwart and RV Seward Johnson as well as to OceanPact for the sampling activities. We are deeply grateful for the invaluable guidance and support provided by Rosa Carvalho Gamba, your legacy wisdom continues to inspire us even in her absence. A special thanks to the editorial board and anonymous reviewers for their careful reading of our manuscript and their many insightful comments and suggestions.

## Author contributions

Julio Cezar Fornazier Moreira: investigation, methodology, formal analysis, visualization, writing - original draft, Writing - review & editing.

Fluvio Modolon: investigation, methodology, formal analysis, visualization, writing - original draft, Writing - review & editing.

Natascha M. Bergo: investigation, methodology, visualization, writing - original draft, Writing - review & editing.

Fabiana S. Paula: investigation, methodology, writing - review & editing. Mateus Gustavo Chuqui: methodology, visualization, formal analysis.

Frederico P. Brandini: project administration, writing - review & editing.

Daniel L. Moreira: funding acquisition, project administration, writing - review & editing.

Celio Roberto Jonck: funding acquisition, project administration, writing - review & editing.

Vivian H. Pellizari: supervision, writing - original draft, Writing - review & editing.

## SUPPLEMENTARY MATERIAL

**Summary**

*Supplementary Data*

**Sup Data 1.** Spreadsheet containing the processed input data of abundance matrix at family level.

**Sup Data 2.** Spreadsheet containing the complete metadata information.

*Supplementary Table*

**Table Sup 1.** Quality measures from SOM.

*Supplementary Figures*

**Fig Sup 1.** HeatMap 2019

**Fig Sup 2.** HeatMap 2021

**Fig Sup 3.** Elbow Method.

**Fig Sup 4.** Indicspecies for SOM’s Association 1.

**Fig Sup 5.** Indicspecies for SOM’s Association 2.

**Fig Sup 6.** Indicspecies for SOM’s Association 3.

**Fig Sup 7.** Indicspecies for SOM’s Association 4.

**Fig Sup 8.** Indicspecies for SOM’s Association 5.

**Fig Sup 9.** Feature Importance of environmental properties for absolute abundance of autotrophic, heterotrophic and total prokaryotic cell measurements.

**Fig S10.** Influence Score of environmental properties for absolute abundance of autotrophic, heterotrophic and total prokaryotic cell measurements.

