## Supplementary Information for "A Machine Learning Approach Elucidates Spatial Patterns of Environmental Properties Driving Microbial Composition Over Santos Basin, South Atlantic"

[4] PETROBRAS Research Center. Centro de Pesquisas Leopoldo Américo Miguez de Mello (CENPES), Rio de Janeiro, Brazil

**Corresponding author**

These authors contributed equally to this work

**Summary**

*Supplementary Table*
**Table Sup 1.** Quality measures from SOM.

*Supplementary Figures*
**Fig Sup 1.** HeatMap 2019
**Fig Sup 2.** HeatMap 2021
**Fig Sup 3.** Elbow Method.
**Fig Sup 4.** Indicspecies for SOM's Association 1.
**Fig Sup 5.** Indicspecies for SOM's Association 2.
**Fig Sup 6.** Indicspecies for SOM's Association 3.
**Fig Sup 7.** Indicspecies for SOM's Association 4.
**Fig Sup 8.** Indicspecies for SOM's Association 5.
**Fig Sup 9.** Feature Importance of environmental properties for absolute abundance of autotrophic, heterotrophic and total prokaryotic cell measurements.
**Fig S10.** Influence Score of environmental properties for absolute abundance of autotrophic, heterotrophic and total prokaryotic cell measurements.

**Table Sup 1.** Quality measures from SOM analysis. The measurements were calculated using aweSOM R package (v1.3).

| Layer: | Coords.Depth_scaled | taxa_total_sqrt4 | mean |
|---|---|---|---|
| Quantization | 0.683809615 | 0.50749 | 0.595648 |
| Explain.var | 77.14 | 68.66 | 72.9 |
| Topograpahic | 0.800546448 | 0.04645 | 0.423497 |
| Neuron utilization | 0.08 | 0.08 | 0.08 |

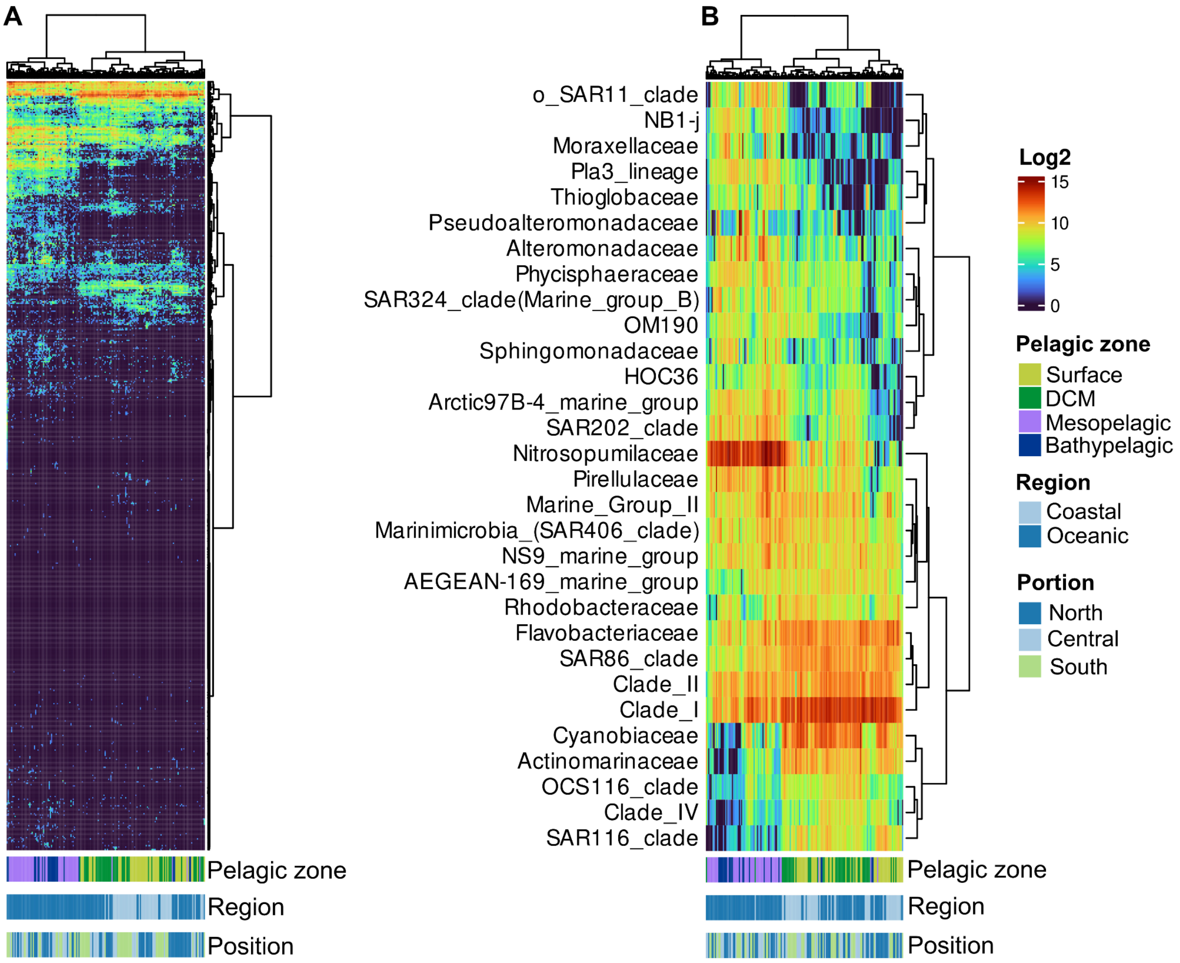


**Figure S1.** Abundance of the top abundance families across the environmental parameters at sampling from 2019. Panel "A" shows the entire microbiome from Santos Basin whilst panel "B" shows the top 30 most abundant taxa.

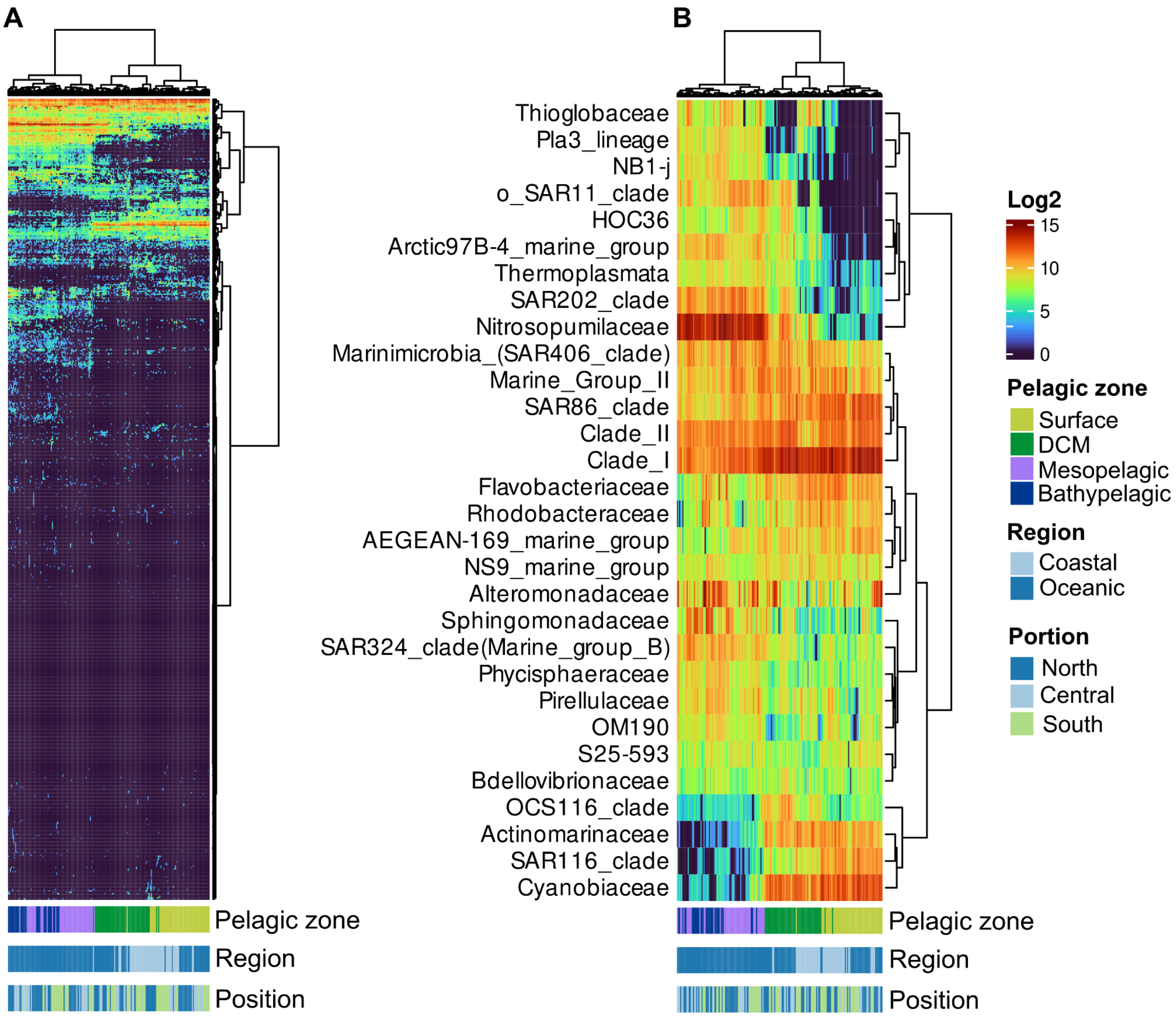


**Figure S2.** Abundance of the top abundance families across the environmental parameters at sampling from 2021. Panel "A" shows the entire microbiome from Santos Basin whilst panel "B" shows the top 30 most abundant taxa.

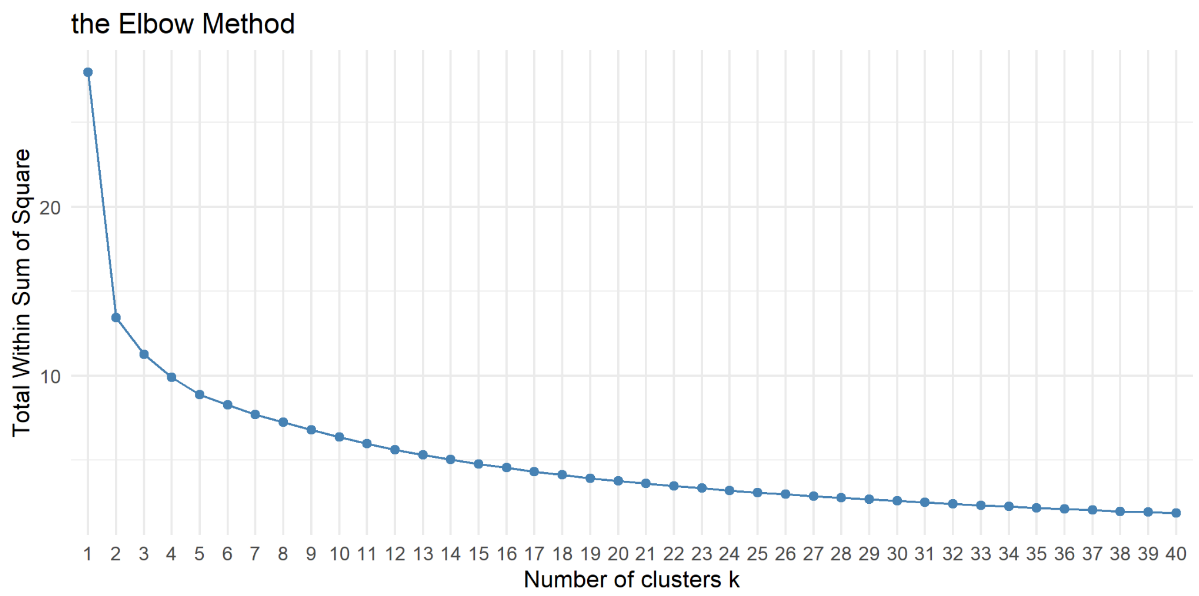


**Figure S3**. Elbow method to determine the optimal number of clusters for hierarchical clustering based on the codebooks generated by Self-Organizing Maps (SOM). The plot shows the total within-cluster sum of squares (WSS) as a function of the number of clusters *k*.

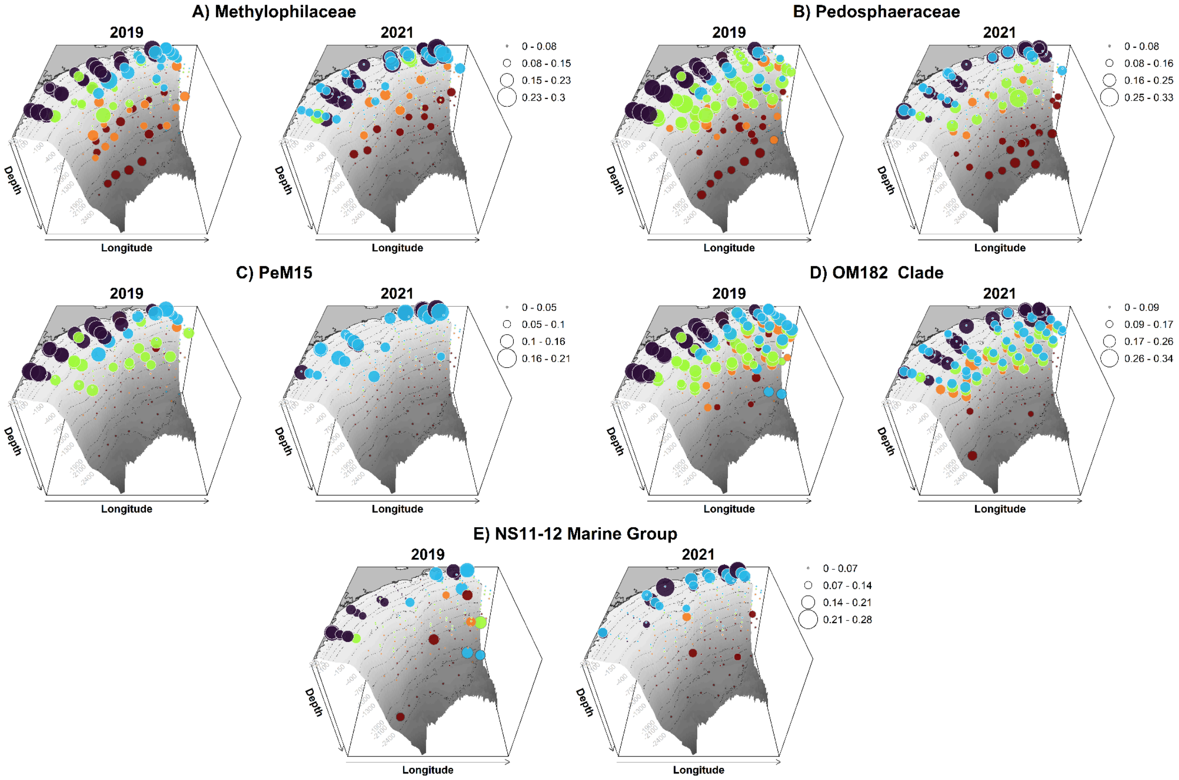


**Fig Sup 4.** Indicator species of SOM Association 1: Relative abundance and distribution patterns of the top indicator microbial families in SOM Association 1 across the Santos Basin. Black circles highlight SOM's Association 1.

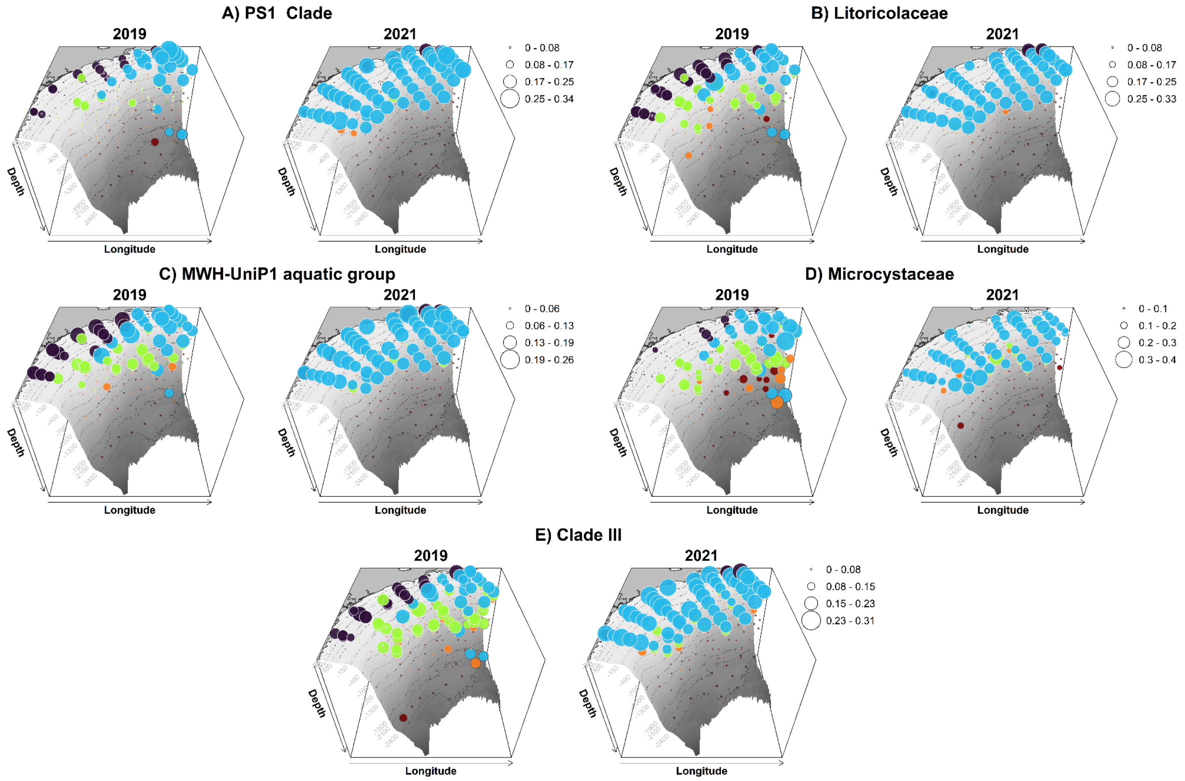


**Fig Sup 5.** Indicator species of SOM Association 2: Relative abundance and distribution patterns of the top indicator microbial families in SOM Association 2 across the Santos Basin. Blue circles highlight SOM's Association 2.

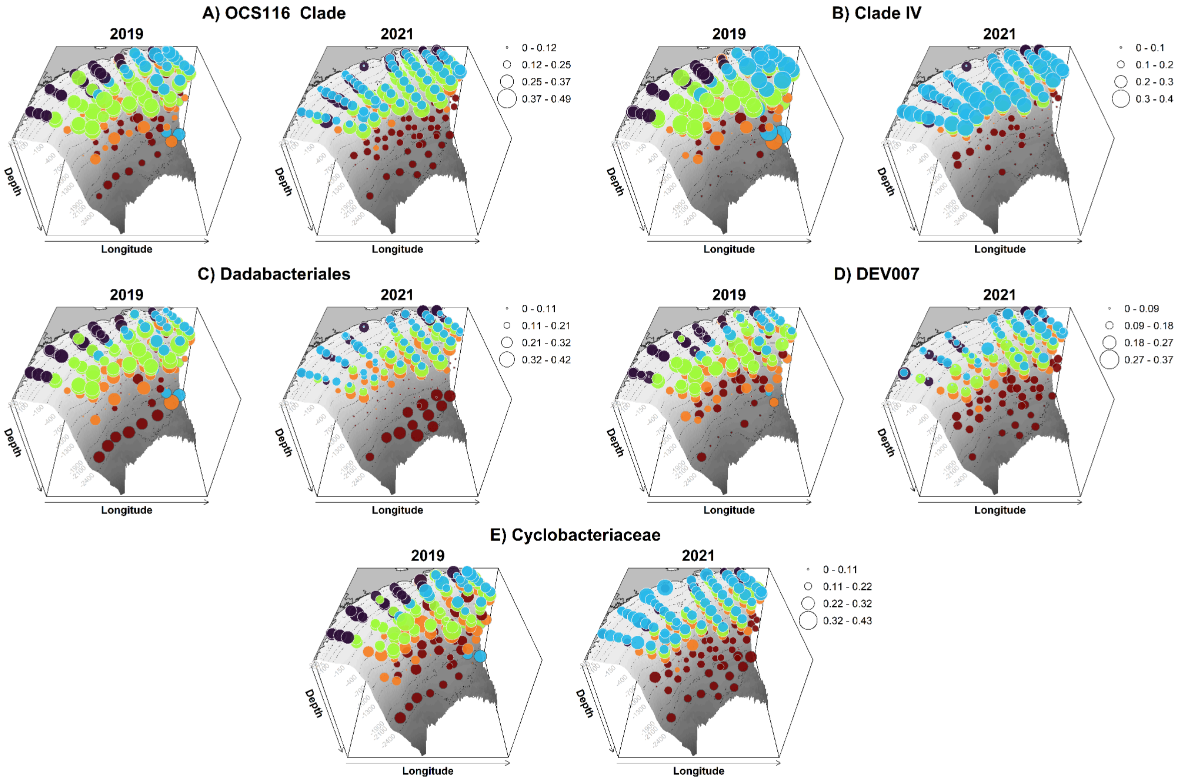


**Fig Sup 6.** Indicator species of SOM Association 3: Relative abundance and distribution patterns of the top indicator microbial families in SOM Association 3 across the Santos Basin. Green circles highlight SOM's Association 3.

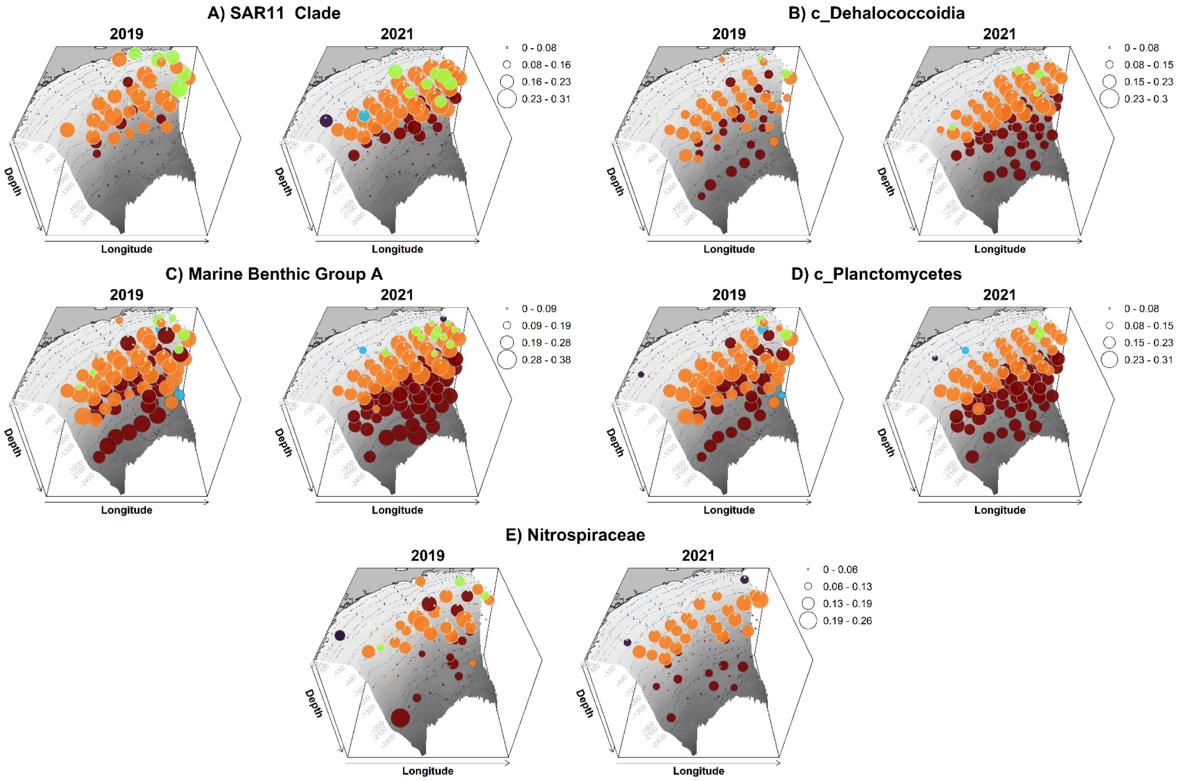


**Fig Sup 7.** Indicator species of SOM Association 4: Relative abundance and distribution patterns of the top indicator microbial families in SOM Association 4 across the Santos Basin. Orange circles highlight SOM's Association 4.

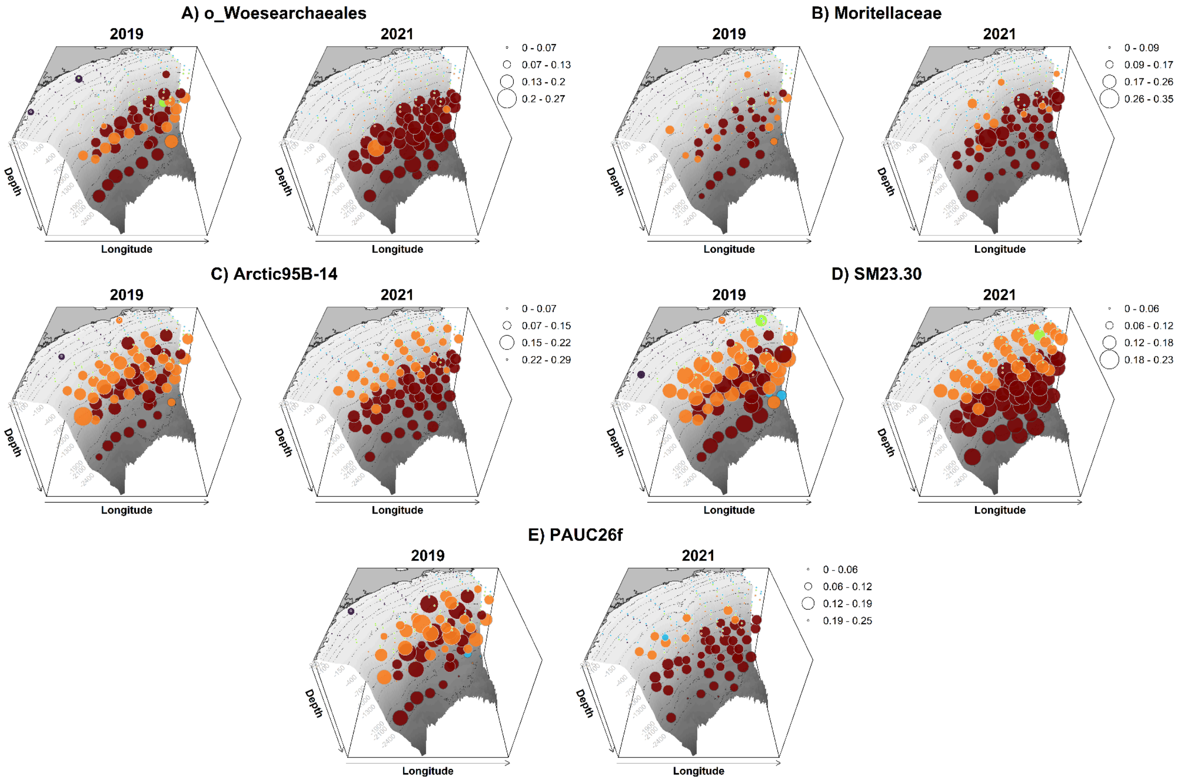


**Fig Sup 8.** Indicator species of SOM Association 5: Relative abundance and distribution patterns of the top indicator microbial families in SOM Association 5 across the Santos Basin. Red circles highlight SOM's Association 5.

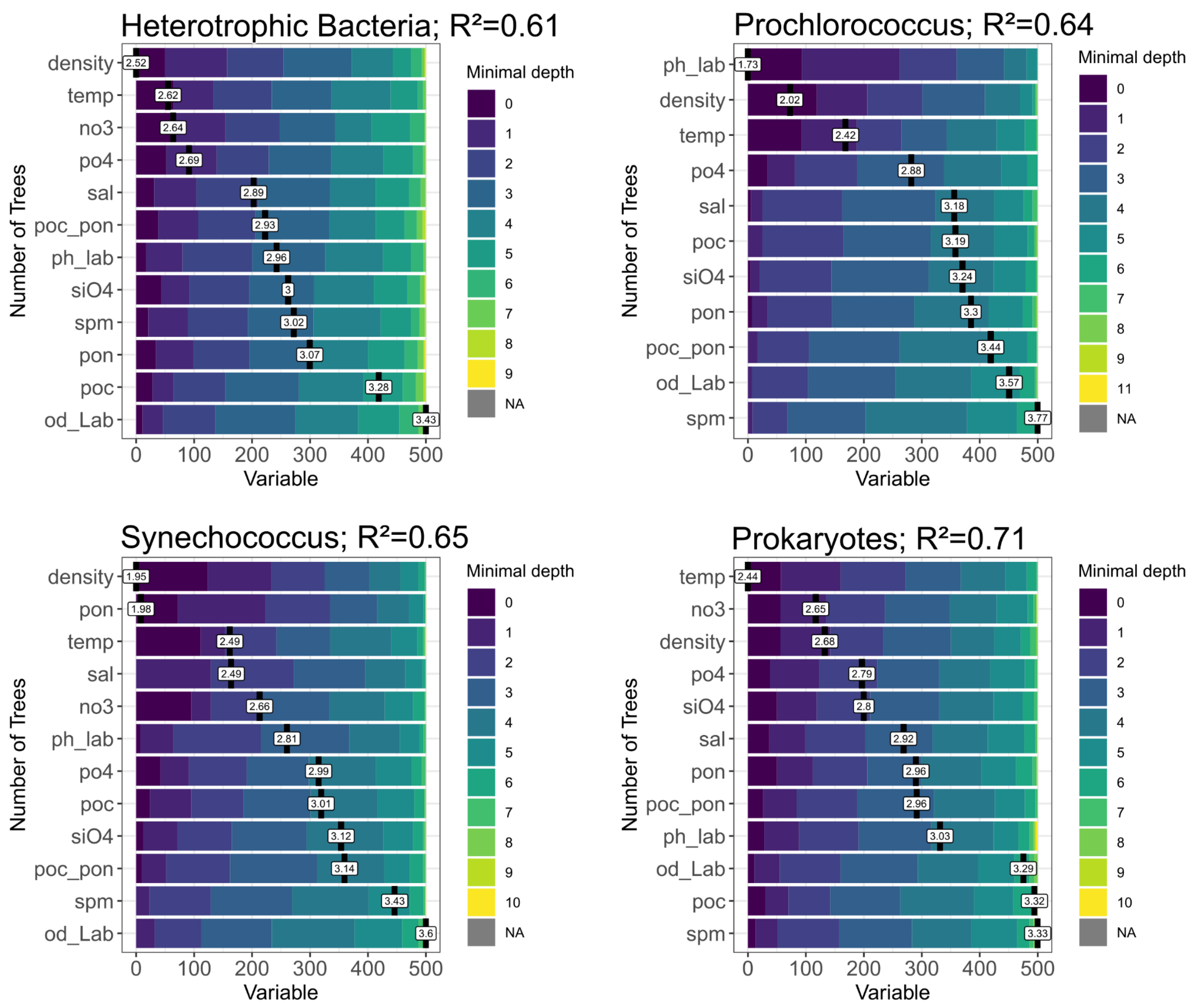


**Fig S9.** Feature Importance of environmental properties for absolute abundance of autotrophic, heterotrophic and total prokaryotic cell measurements.

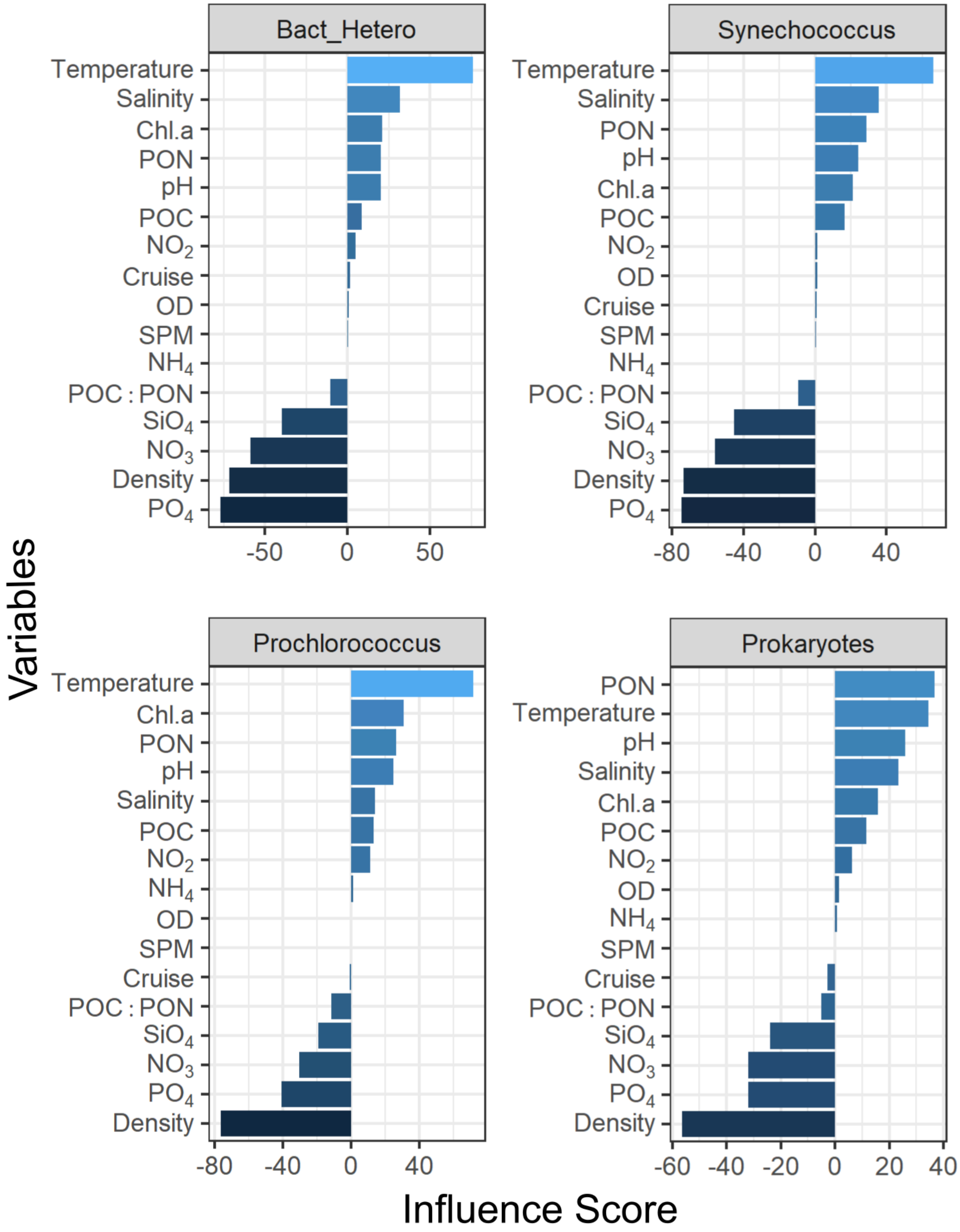


**Fig S10.** Influence Score of environmental properties for absolute abundance of autotrophic, heterotrophic and total prokaryotic cell measurements.
